# Density-dependent effects of mortality on the optimal body size of a habitat shift: Why smaller is better despite increased mortality risk?

**DOI:** 10.1101/728956

**Authors:** P. Catalina Chaparro-Pedraza, André M. de Roos

**Affiliations:** Institute for Biodiversity and Ecosystem Dynamics, University of Amsterdam, 1090 GE Amsterdam, the Netherlands

**Keywords:** size-dependent mortality, size-selective mortality, optimal life history trait, habitat shift

## Abstract

Many animal species across different taxa change their habitat during their development. An ontogenetic habitat shift enables the development of early vulnerable-to-predation stages in a safe ‘nursery’ habitat with reduced predation mortality, while less vulnerable stages can exploit a more risky, rich feeding habitat. Therefore, the timing of the habitat shift is crucial for individual fitness. We investigate the effect that size-selectivity in mortality in the rich feeding habitat has on the optimal timing of the habitat shift using a population model and the adaptive dynamics approach. We show that the size-selective nature of mortality in this habitat affects density-dependent body growth rate in the nursery habitat and thus the optimal timing of the habitat shift. This is caused by the effect exerted by size-dependent mortality on the size distribution of the population that results in strong competition in the nursery habitat. We furthermore find that, as a consequence of this effect, increased size-selectivity in mortality in the rich feeding habitat causes the optimal body size to shift habitat to decrease. Our results reveal the interdependence between population structure and life history traits, and highlight the need for integrating ecological interactions in the study of the evolution of life histories.

## Introduction

Interactions between organisms do not remain constant throughout their lives. Instead, the outcome of encounters between competitors, between prey and predator or between parasite and host depends on the developmental stage of the interacting organisms. An increase in body size is the most important ecological aspect of ontogenetic development as it determines to a large extent those interactions besides feeding, growth and reproduction (de Roos and Persson 2013). Ecological interactions, therefore, change with the increase in size during ontogeny. In particular, smaller or younger individuals of diverse fish (Sogard 1997; Krause et al. 1998; Hampton 2000), amphibian (Arendt, 2009; Rudolf, 2018; Semlitsch, 1990), reptile (Ferguson and Fox 1984; Keren-Rotem et al. 2006) and invertebrate species (Rudolf and Armstrong 2008; Boulton and Polis 1999; Keller and Ribi 1993) experience higher predation or cannibalistic risk than larger ones. To reduce the risk of injury or lethal interactions, small individuals often avoid areas with predators or larger conspecifics (Ohgushi et al. 2012) by using the same habitat differentially (Diehl and Eklov 1995) or using two different habitats for small and large individuals (Dodson et al. 2009).

Among vertebrate animal species, ontogenetic habitat shifts, understood as the use of different habitats in different stages of the life history, have been documented in a wide range of fish and amphibian species, as well as, in some reptiles (Werner and Gilliam 1984; Keren-Rotem et al. 2006). By switching habitat small individuals can develop in a safe ‘nursery’ habitat with reduced predation mortality, while large individuals exploit a riskier, but richer feeding habitat. For instance, salmon and other anadromous species utilize freshwater streams as breeding habitats that offer a reduced predation mortality for embryos and larval stages compared to their marine counterparts, whilst large less vulnerable-to-predation individuals exploit the productive marine grounds at high latitudes (Dodson et al. 2009). Likewise, in several fish species associated with coral reefs, only large individuals are actually present in this highly productive habitat, and small vulnerable-to-predation stages occur in habitats with lower predation mortality such as mangroves and seagrass beds (Cocheret De La Morinière et al. 2002; Kimirei et al. 2013).

The physical separation between different ontogenetic habitats implies multiple changes in ecological conditions experienced by individuals during the habitat shift: the individuals do not only experience a change in predation vulnerability during the habitat shift, but also a diet shift (Hobson 1999), as well as changes in intraspecific competition and food abundance (Diehl and Eklov 1995; Keren-Rotem et al. 2006). Generally, the ‘nursery’ habitat is of relatively small size and low productivity compared to the habitat occupied by older individuals. As a consequence, individuals in the former experience increased density while density-dependence is very low to negligible in the latter (Diehl and Eklov 1995; Jonsson et al. 1998; Cocheret De La Morinière et al. 2002). This relaxation of the intraspecific competition in the habitat occupied by older individuals leads to an increase in food abundance after the habitat shift that, in turn, increases the available energy to allocate to both somatic growth and reproduction. An early habitat shift thus enables an early onset of rapid growth and reproduction. However, small individuals are more vulnerable to predation on arrival in the second habitat. A late habitat shift would allow them to reach a larger body size before entering the riskier habitat and therefore lowers predation mortality at the expense of an extended period of slow growth in the first habitat. Under such a trade-off, the timing to shift habitat is a crucial determinant of individual fitness.

Werner and Gilliam (1984) concluded that when the two habitats differ in size-specific growth and mortality rates (indicated with *g* and *μ*, respectively), fitness is maximized when the switching size minimizes the ratio of mortality to growth rate (also referred to as the “*μ/g* rule”). However, this conclusion is based on an individual optimization in an invariant environment. It therefore ignores density-dependent processes at the population level caused by the interactions among individuals, such as the difference in intraspecific competition in the two habitats mentioned above. Furthermore, the optimal size to switch habitats determines the outflow and inflow of individuals in the two habitats through growth and reproduction and thus the densities of individuals in each habitat. These changes in population density affect intraspecific competition that, in turn, affects individual growth rate and therefore the optimal strategy to switch habitats. A few studies have investigated the optimal timing of a habitat or niche shift incorporating intraspecific competition (Claessen and Dieckmann 2002), but the role of mortality in the rich feeding habitat and its link with body size has not been explored yet.

Although size-dependent mortality due to predation is usually the main mortality source in the rich feeding habitat, physical mortality factors that cause random mortality across all size classes (i.e. uniform mortality), such as oxygen depletion and temperature extremes, can sometimes override size-dependent mortality (Sogard 1997). In this study, we investigate how size-selectivity in mortality in the rich feeding habitat affects the optimal timing of a habitat shift. To do so, we use a size-structured population model for a consumer-resource interaction that incorporates food-dependent individual growth for the consumers. We analyze the ecological dynamics predicted by the model and use the adaptive-dynamics approach to determine the optimal body size of the habitat shift in the context of these ecological dynamics.

## Methods

### The model

We formulate a model that accounts for a population in two habitats. We assume that in each habitat the individuals exploit a different resource. The population is structured by individual body size. Individual resource consumption, somatic growth, survival and reproduction are following continuous-time dynamics. We study the population in the ecological equilibrium state.

### Life cycle

Individuals are born in the ‘nursery’ habitat (hereafter habitat 1) with size *l*_0_ where they remain until they reach a body size *l_s_* when they shift to the rich feeding habitat (occupied by older individuals, hereafter habitat 2). Juvenile individuals mature and start to reproduce at a body size *l_m_*.

### Habitats

Density-dependence due to competition for food is considered to be strong in habitat 1, directly influencing growth in body size like in salmonids (Walters et al. 2013), so we assume the food or resource density in this habitat to be depleted by the foraging of consumer individuals. In the absence of consumers, the resource is assumed to follow a semi-chemostat growth dynamics with maximum density *R*_1 max_ and growth rate *ρ* (for an explanation and justification of this type of growth dynamics, see Persson et al. (1998)). Dynamics of the resource density *R*_1_ in habitat 1 in the absence of consumers is hence given by:

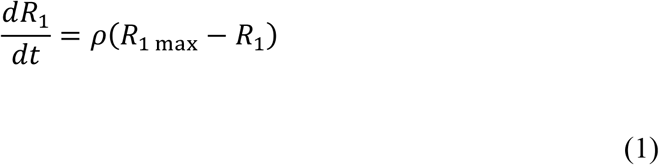

In contrast, in habitat 2, density-dependence is considered negligible, therefore we assume a constant resource density.

### Individual dynamics

The core part of the model is the description of the individual behavior, that is, feeding, growth, reproduction and mortality as a function of the individual state (i.e. body size) and the state of the environment (food availability). In the following sections we describe the individual level dynamics.

- *Feeding* In habitat 1, individuals are assumed to feed on the resource following a Holling type II functional response. So their feeding level *f*_1_ (or scaled functional response) is given by:

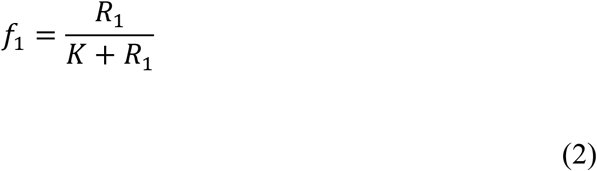

where *K* is the half-saturation resource density. In habitat 2 individuals feed at a constant feeding level *f*_2_.
- *Dynamic energy budget model: Individual state and fecundity* The model follows the bioenergetics approach introduced by Kooijman and colleagues (Nisbet et al. 2000; Kooijman 2010; Kooijman and Metz 1984) in which the energy allocation to somatic and reproductive metabolism is proportional to a fraction *κ* and a *1−κ* of the total energy assimilation rate, respectively. More specifically, we adopt the model developed and described in detail by Jager et al. (2013). Below we provide only a concise synopsis of the model. Individuals are characterized by the state variable structural mass, *W*. Body length *l* and structural mass *W* are related to each other following Jager et al. (2013):

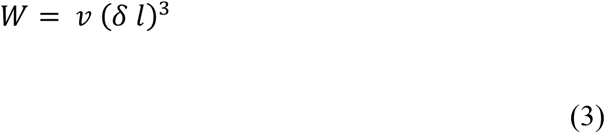

where *ν* is the density of structural mass and *δ* is a shape coefficient factor. Hereafter we refer to *l* as body size. The acquisition and utilization of energy are described by equations (4) to (10). The energy assimilation rate is given by:

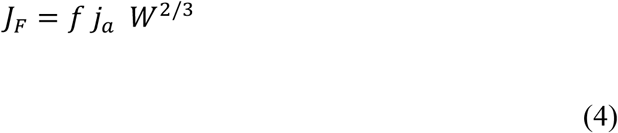

where *f* is the feeding level in either habitat, *j_a_* is the maximum area-specific assimilation rate and the surface area for assimilation is assumed to scale with structural mass to the power of 2/3. Metabolic maintenance costs are the product of the mass-specific maintenance costs *j_m_* and the structural mass:

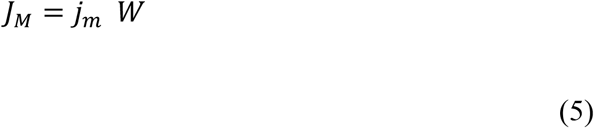 Assimilates are assumed to split into two energy fluxes: the *κ* flux and the 1 − *κ* flux. The *κ* flux is first used to cover metabolic maintenance costs, while the remaining flux *J_W_* is used to synthesize structural mass. On the other hand, the 1 − *κ* flux *J_S_* is allocated to reproduction when adult and to maturation when juvenile (Jager et al. 2013). Hence,

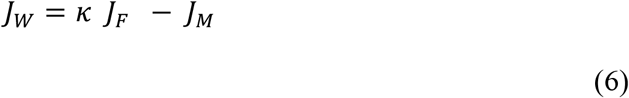

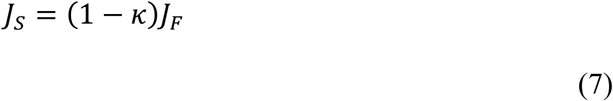 Therefore, the dynamics of structural mass *W* as a function of age *a* is described by the differential equation:

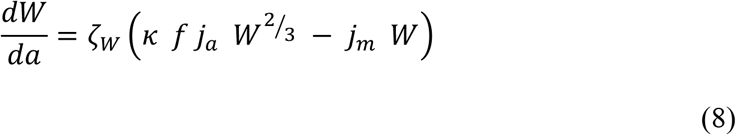 The parameter *ζ_W_* in equation (8) represents the efficiency with which assimilates are converted into structural mass. The previous equation can be rewritten as growth in body size *γ*(*f, l*) by substituting structural mass by body size (from equation 3) and doing some manipulations:

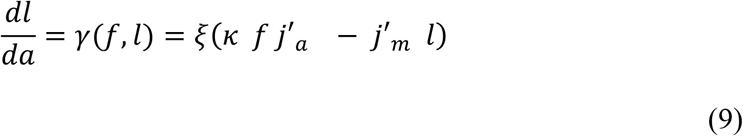 The compound parameter *ξ* characterizes the growth rate in size and is defined as *ζ_W_*/(3 *ν δ*^3^). While the compound parameters *j′_a_* and *j′_m_* correspond to the assimilation and maintenance rates with respect to body length and are defined as *j_a_ ν*^2/3^ *δ*^2^ and *j_m_ ν δ*^3^ respectively. Reproduction is assumed continuous. Adult fecundity after the substitution of structural mass by body size is described by:

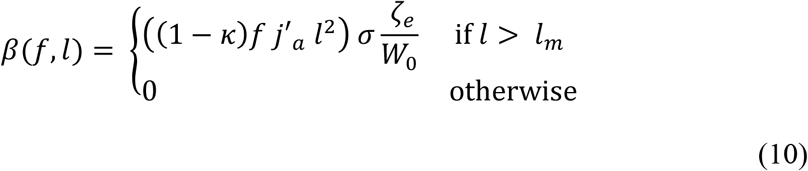 The number of offspring produced per unit time is dependent on the yield for the conversion of assimilates into eggs *ζ_e_* and the newborn structural mass *W*_0_ = *ν* (*δ l_o_*)^3^. In addition, although we do not consider explicitly the egg stage, we do consider the egg survival *σ* that reduces the number of individuals effectively recruited as newborns. We study the research question only in the ecological equilibrium state when the condition *κ f j′_a_* > *j′_m_ l* is fulfilled, therefore individuals do not starve and cannot shrink in size.
- *Survival* Individuals in habitat 1 may die from background mortality *μ*_1_ and in habitat 2 from either background *μ*_2 *b*_ or predation mortality *μ*_2 *p*_. Background mortality is assumed to be size-independent and predation mortality is assumed size-dependent. To describe the size-dependent mortality experienced by individuals in habitat 2 we adopt a continuous piecewise-differentiable sigmoid function of body size (fig. 1):

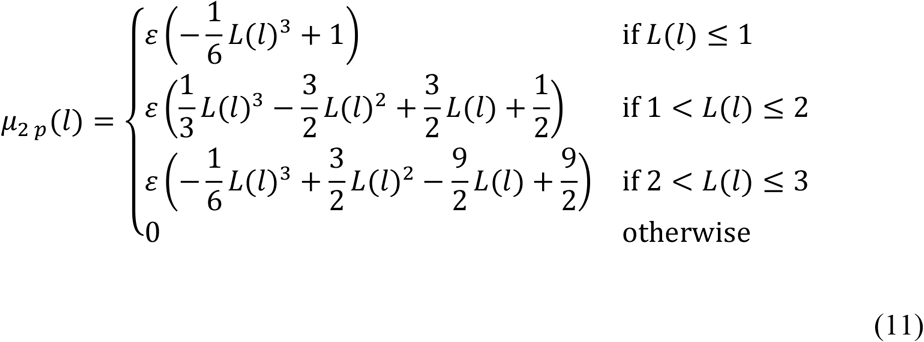

where *L*(*l*) is a scaled body size value, defined as *L*(*l*) = 3 *l/l_ν_*. The sigmoid function is bounded by the maximum size-dependent mortality *ε*, which occurs at *l* = 0, and the maximum vulnerable-to-predation body size *l_ν_* at which size-dependent mortality vanishes. This function has been chosen because the parameters *ε* and *l_ν_* facilitate biological interpretation. However, size-dependent mortality has been commonly described as an exponential function of body size (fig. S1; Gislason et al. 2010; Jørgensen and Holt 2013):

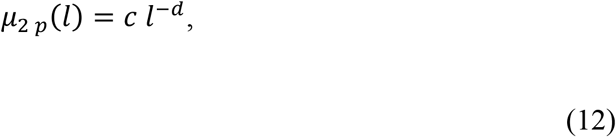

therefore, we test the robustness of our results under this assumption (see Supp. Info.).

**Figure 1.**
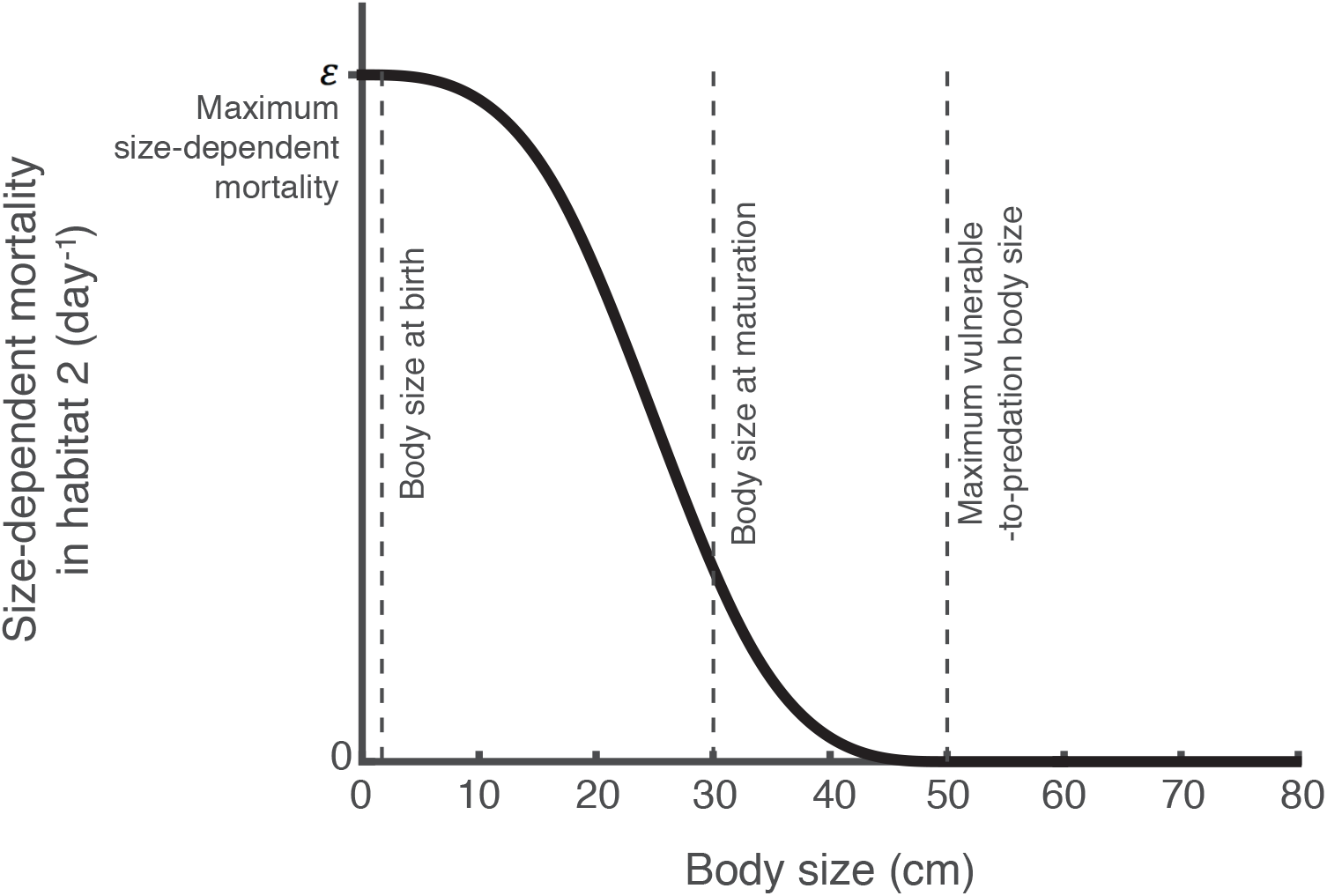
Size-dependent mortality in habitat 2.

The total per capita death rate in habitat 2 *μ*_2_ is the sum of the background and size-dependent mortality.

### Population dynamics

Based on the individual life history described above, the size-structured population model can be formulated by bookkeeping following de Roos et al. (1990) and de Roos (1997). A detailed description of the size-structured population model can be found in the supplementary information.

### Evolutionary dynamics

In this study, we are interested in understanding how size-selectivity in mortality in the rich feeding habitat affects the optimal timing of a habitat shift. We study the evolution of the body size at the habitat shift *l_s_* using the adaptive dynamics framework, a suitable framework to analyze phenotypic evolution. Specifically, we consider a population that evolves through the fixation of small and rare mutations in this trait while otherwise being monomorphic (Geritz et al. 1998). A mutation is eventually fixed if the population growth rate of mutant individuals in the environment imposed by the resident population, which, in our model, is the food resource density in the habitat 1 (*R*_1_), is positive. This population growth rate represents the mutant’s invasion fitness, which for the resident equals 0 when it is at an equilibrium state. The fitness of a mutant therefore depends on its own strategy and on the strategy of the resident. We use the lifetime reproductive output *R*_0_ as a measure of invasion fitness following Mylius and Diekmann (1995). A mutant can invade if its lifetime reproductive output *R*_0_(*l′_s_*|*l_s_*) > 1; where *l′_s_* is its own strategy, and *l_s_* is the strategy of the resident population. When a mutant phenotype invades, it spreads and the new population reaches the ecological attractor. Then, it can be invaded by another mutant that has an invasion fitness larger than one. In this way,the population experiences a succession of mutations and evolves in the direction of the selection gradient

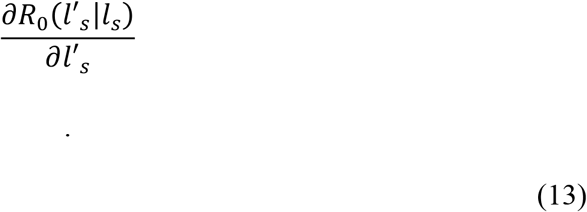

The point where the selection gradient becomes 0 is the Evolutionarily Singular Strategy (ESS), which can be evolutionary stable or unstable. In the first case, no other mutant can invade in the population and the ESS is a continuously stable strategy (CSS), while in the latter evolutionary branching can occur (Geritz et al. 1998).

The lifetime reproductive output *R*_0_ is numerically calculated and used in the canonical equation of adaptive-dynamics theory (Dieckmann and Law 1996; Durinx et al. 2008) to determine the resultant evolutionary trajectories. A detailed description of the evolutionary model and the canonical equation can be found in the supplementary information.

### Model parameterization

All biomass densities are expressed in grams per liter, and time in the ecological timescale is expressed in days. Parameters of the population with a habitat shift are loosely based on the biology of Atlantic salmon (*Salmo salar*). Scaling coefficients of assimilation and metabolic rate (*j_a_, j_m_*) were estimated from regressions of Koskela, Pirhonen, and Jobling (1997) with the method of Jager, Martin, and Zimmer (2013) and corrected for a temperature of 10 °C (although there is large variation in temperature in wild populations, FishBase lists an approximate preferred temperature of 9.3 °C for Atlantic salmon (Froese and Pauly 2018)). The yield parameter of assimilates from ingested food efficiency has only a scaling effect on the population biomass equal for all stages, therefore, as it does not have any qualitative effect on the results, we assume a value of 1 g/g. The shape coefficient was estimated from regressions of Sutton, Bult, and Haedrich (2000); its estimated value is similar to shape coefficient by Pecquerie et al. (2011) for Pacific salmon species. Other parameters of the bioenergetics model are taken from Jager et al. (2013), while life history traits (i.e. body size at birth) were derived from reported data in the literature (table S1). Atlantic salmon are considered mature when they return to the streams to spawn (around 50 cm; Hutchings and Jones 1998), however at this point, individuals had already accumulated large amounts of energy for reproduction. It is unknown, however, when they start to allocate this energy to reproduction. Since we assume reproduction to be a continuous process in the model (i.e. energy allocated to reproduction is immediately converted into offspring), we chose a threshold for maturation lower than the body size at which Atlantic salmon has been documented to return (30 cm).

The model always approaches a stable ecological equilibrium for the parameterization considered. Therefore, the right-hand side of equation 9 is always positive, and possible starvation conditions of the consumers can be ignored.

### Model analysis

We focus on populations in an ecological equilibrium and use the PSPManalysis software package (de Roos, 2018) to numerically compute and continue the equilibrium of the size-structured population model described above as a function of model parameters. We perform the evolutionary analysis using the PSPManalysis package as well, as it allows us to detect, classify and continue an ESS as a function of *ε*, the scaling factor in the size-dependent mortality function *μ*_2 *p*_, and *μ*_2 *b*_, the size-independent mortality parameter in habitat 2.

We are interested in understanding how size-selectivity in mortality in the rich feeding habitat (habitat 2) affects the optimal timing of a habitat shift. However, a simple increase in size-dependent mortality does not only increase the size-selective nature of mortality but also the total mortality experienced in this habitat. Hence, the effect of the size-selectivity of mortality per se can only be unraveled while maintaining total mortality constant in the habitat 2. To evaluate evolutionary responses we therefore follow a specific approach, in which the contribution from size-dependent mortality sources is increased but overall mortality in habitat 2 is kept constant through a simultaneous decrease in size-independent mortality. More specifically, we find the ESS value for the body size to shift habitat when there is only size-independent mortality *μ*_2 *b*_ equal to 0.006 day^-1^ in habitat 2 and adopt this as our starting, reference population (body size at habitat shift = 19.5 cm). A size-independent mortality *μ*_2 *b*_ equal to 0.006 day^-1^ implies that an individual has an expected lifetime of 167 days in habitat 2 from the moment it shifts habitats. Adopting this ESS body size for the resident phenotype in case it only experiences size-independent mortality *μ*_2 *b*_ of 0.006 day^-1^, we infer two combinations of size-independent mortality *μ*_2 *b*_ and maximum size-dependent mortality *ε* that also result in a life expectancy of 167 days in habitat 2 (Pop. # 2 and 3 in table 1). To infer these combinations we numerically compute an individual’s life expectancy after entering habitat 2 by integrating the following differential equation for its survival *S*(*τ*) as a function of the time *τ* it has spent in habitat 2:

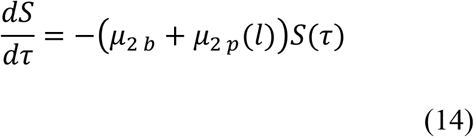

**Table 1.** Mortality parameters used in Fig. 2, 3 and 4

| Pop. # | Body size at habitat shift<sup>a</sup> (cm) | Life expectancy in habitat 2 (days) | Maximum size-dependent mortality $\varepsilon$ | Contribution of size-dependent mortality <sup>b</sup> | Size-independent mortality $\mu_{2b}$ | Contribution of size-independent mortality <sup>b</sup> |
| --- | --- | --- | --- | --- | --- | --- |
| Increasing size-selectivity in mortality in habitat 2 |  |  |  |  |  |  |
| 1<sup>c</sup> | 19.5 | 167 | 0 | 0% | 0.006 | 100% |
| 2 | 19.5 | 167 | 0.0036 | 33.3% | 0.004 | 66.7% |
| 3 | 19.5 | 167 | 0.0075 | 66.7% | 0.002 | 33.3% |
| Decreasing size-selectivity in mortality in habitat 2 |  |  |  |  |  |  |
| 4<sup>c</sup> | 21.4 | 144 | 0.015 | 98.3% | 0.0001 | 1.7% |
| 5 | 21.4 | 144 | 0.01 | 70.1% | 0.0021 | 29.9% |
| 6 | 21.4 | 144 | 0.005 | 36.7% | 0.0044 | 63.3% |

<sup>a</sup> Body size at habitat shift is ESS before change in size-selectivity in mortality.
<sup>b</sup> Contribution of each source of mortality to total mortality in habitat 2.
<sup>c</sup> Starting, reference population

while simultaneously integrating the differential equation (9) for its growth in body size *l*.

We subsequently start with a resident population that is characterized by the ESS value for body size to shift habitat while experiencing a size-independent mortality equal to 0.006 day^-1^ in habitat 2 and assess its evolutionary response when the mortality changes to one of the inferred combinations of size-independent mortality *μ*_2 *b*_ and maximum size-dependent mortality *ε* that also lead to an average life expectancy of 167 days in habitat 2 (fig. 2A). We perform an analogous analysis while decreasing the size-selectivity in mortality in habitat 2: We adopt as our starting, reference population, one characterized by the ESS value for the body size to shift habitat when size-dependent mortality is the main source of mortality with a maximum value *ε* of 0.015 day^-1^ in habitat 2 (body size to shift habitat = 21.4 cm; a very low size-independent mortality equal to 0.0001 day^-1^ in habitat 2 is introduced to avoid that individuals with a body size larger than the maximum vulnerable-to-predation body size are immortal). We identify again two other combinations of maximum size-dependent mortality *ε* and size-independent mortality *μ*_2 *b*_ that result in the same life expectancy in the habitat 2 (Pop. # 5 and 6 in table 1) as the size-dependent mortality with maximum value *ε* of 0.015 day^-1^ for individuals shifting habitat at the ESS body size and study the evolutionary response of the starting resident population to a change to the two inferred mortality schedules (fig. 2B). This procedure ensures that the evolutionary change we observe is strictly due to the variation in size-selectivity in mortality and not to a change in total mortality.

**Figure 2.**
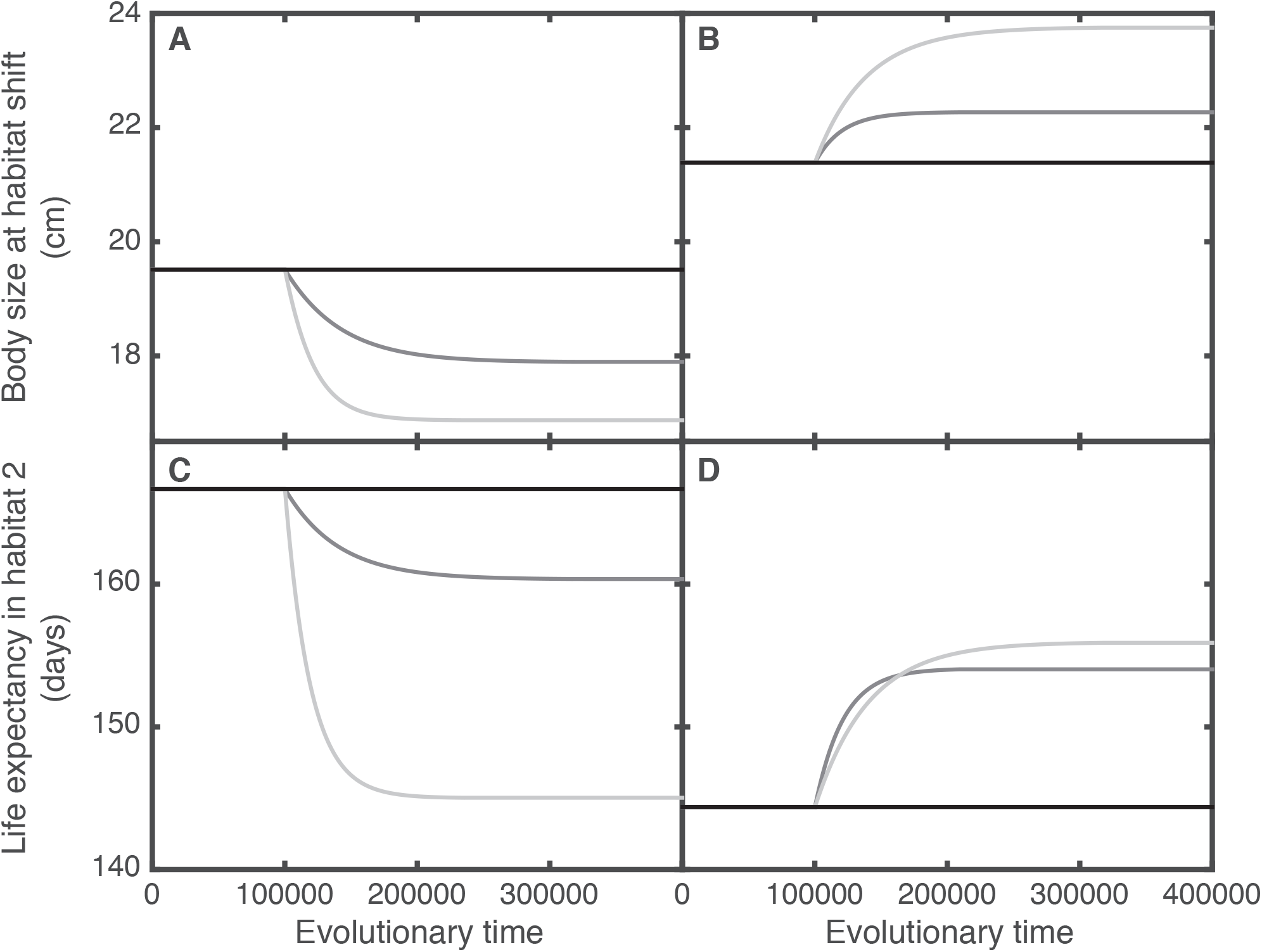
Effects of increased (A, C) and decreased (B, D) size-selectivity in mortality in habitat 2 on the optimal timing of the habitat shift (A, B) and life expectancy in habitat 2 (C, D). When size-selectivity increases (A, C), evolutionary dynamics starts from the ESS value for the body size at habitat shift (19.5 cm) when individuals experience only size-independent mortality 0.006 day^-1^ until time 100 000. At this time, the change in size-selectivity in mortality in habitat 2 occurs (changes in size-dependent and size-independent mortality according to table 1: Pop. # 1 corresponds to the black line, 2 to the dark grey line and 3 to the light grey line). When size-selectivity decreases (B, D), evolutionary dynamics starts from the ESS value for the body size at habitat shift (21.4 cm) when individuals mainly experience a maximum size-dependent mortality 0.015 day^-1^ until time 100 000. At this time, the change in size-selectivity in mortality in habitat 2 occurs (changes in size-dependent and size-independent mortality according to table 1: Pop. # 4 corresponds to the black line, 5 to the dark grey line and 6 to the light grey line). Other parameter values as in table S1.

For the four combinations of size-dependent and size-independent mortalities mentioned above and shown in Table 1 we use the PSPManalysis software package to compute the evolutionary trajectories of the body size at the habitat shift (evolutionary time 100 000 to 400 000 in fig. 2) as predicted by the canonical equation of adaptive dynamics. We perform an evaluation of the fitness components of the resident and a mutant individual in the environment set by the resident just before and immediately after the change in the size-selectivity of mortality (evolutionary time 100 000) to determine how various fitness components are maximized by selection under the imposed variation in size-selectivity in mortality (fig. 3). Likewise, we assess the effect of the variation in size-selectivity in mortality on the size distribution of the population (fig. 4).

**Figure 3.**
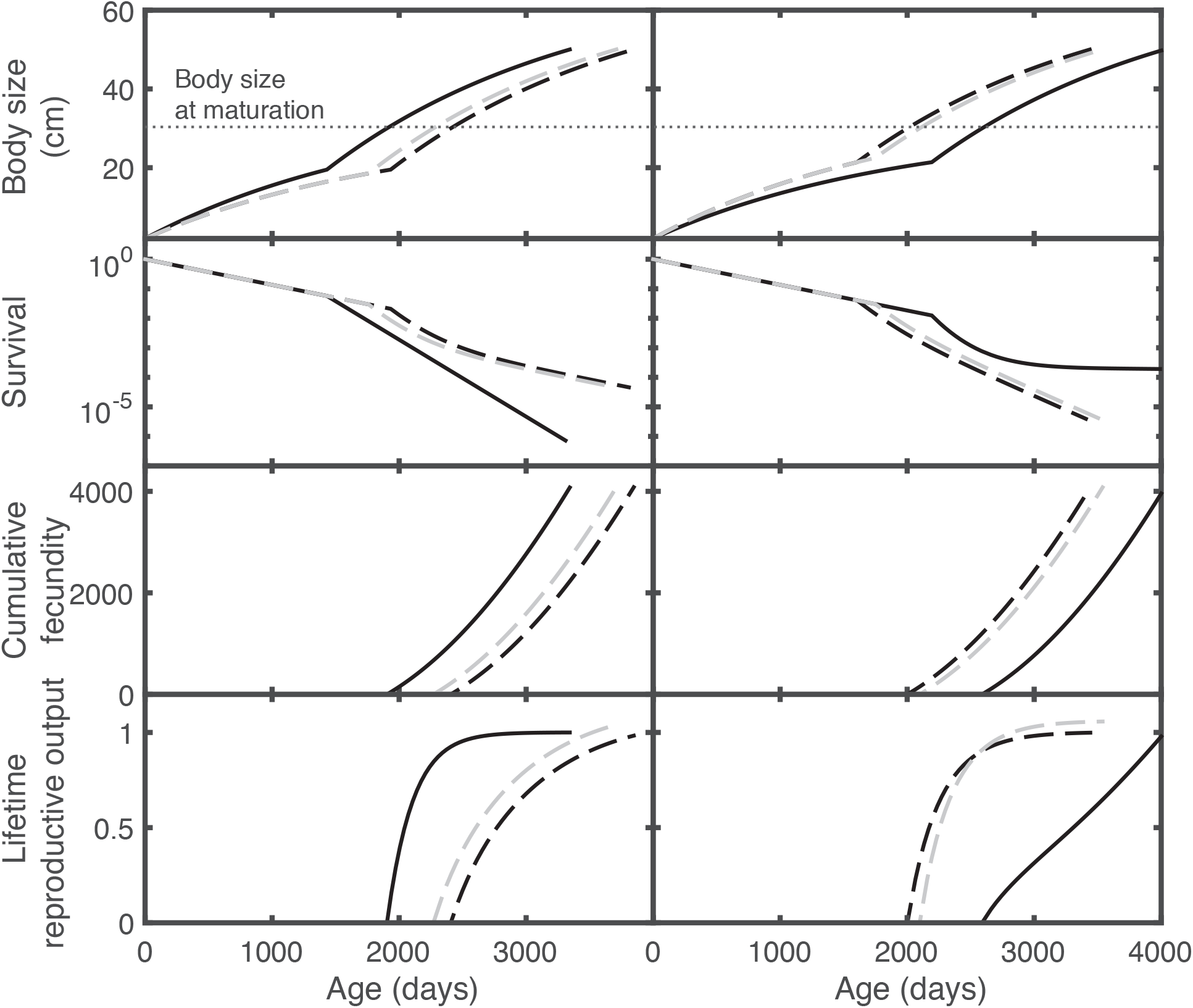
Individual growth, survival, cumulative fecundity and expected lifetime cumulative fecundity of the resident before (black solid lines) and of the resident and mutant phenotypes immediately after (black and rey dashed lines, respectively) size-selectivity in mortality increases (left column) and decreases (right column) in habitat 2. Size-dependent and size-independent mortality parameters before size-selectivity in mortality increases and decreases are as shown in table 1 for Pop. # 1 and 4, respectively; whereas after the increase and decrease they are as shown in table 1 for Pop. # 3 and 6, respectively. Resident phenotypes shift habitat at 19.5 cm (left column) and 21.4 cm (right column) and mutant phenotypes shift habitat at a body size that differs 5% from the body size at habitat shift of the resident phenotype in the direction of the selection gradient (decrease when a smaller body size at habitat shift is selected for in fig. 2, increase if a larger body size at habitat shift is selected for in fig. 2).

**Figure 4.**
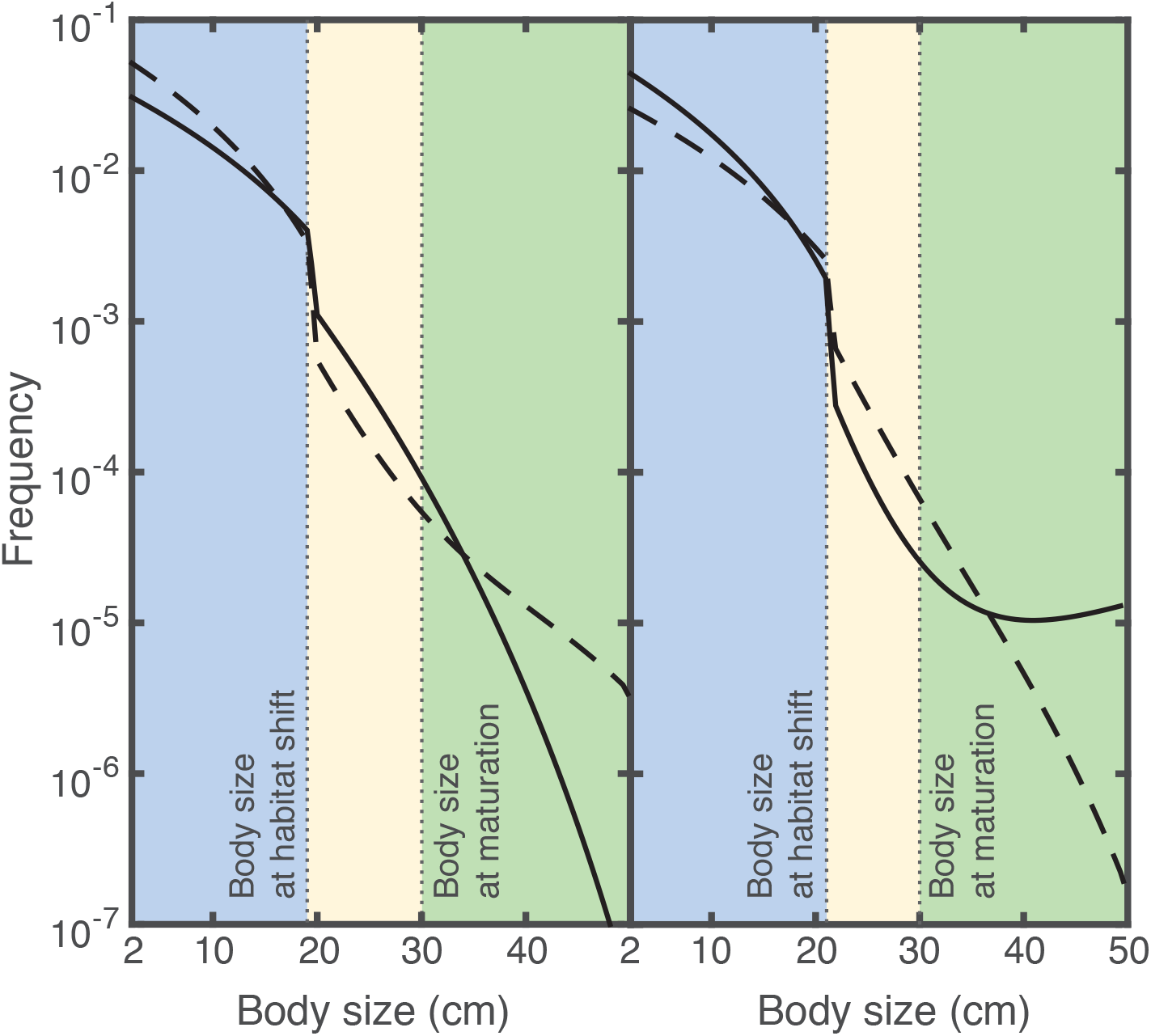
Size-distribution of the resident population just before (solid lines) and immediately after (dashed lines) size-selectivity in mortality increases (left column) and decreases (right column) in habitat 2. Individuals shift habitat at a body size of 19.5 cm (left) and 21.4 cm (right). Juveniles in habitat 1 (blue region) and in habitat 2 (yellow region), and adults (green region) are shown. Size-dependent and size-independent mortality paramenters before size-selectivity in mortality increases and decreases are as shown in table 1 for Pop. # 1 and 4, respectively; whereas after the increase and decrease they are as shown in table 1 for Pop. # 3 and 6, respectively. Other parameter values as in table S1.

A similar procedure is followed to test the robustness of the results under the assumption that size-dependent mortality is an exponential function of the body size. A detailed description can be found in the supplementary information.

## Results

In the first part of this section, we show the evolutionary effects of size-selectivity in mortality in the rich feeding (habitat 2) on the optimal timing of the habitat shift. Subsequently, we present the cause of these evolutionary responses. All evolutionary outcomes described below are continuously stable strategies (CSSs) and therefore locally evolutionarily stable (according to classification by Geritz et al., 1998).

### Increased size-selectivity in mortality in habitat 2 decreases the optimal body size at habitat shift

When size-selectivity in mortality increases in habitat 2 (i.e. the contribution of size-dependent mortality sources to total mortality increases, given a constant life expectancy in habitat 2), the body size at habitat shift decreases, so individuals have an earlier habitat shift with respect to the individuals exposed only to sizeindependent mortality (fig. 2A; notice the same life expectancy at the beginning of the evolutionary trajectories in fig. 2C). As a consequence of the evolution toward a smaller size at habitat shift, life expectancy in habitat 2 decreases as well (fig. 2C).

Conversely, when size-selectivity in mortality in habitat 2 decreases (i.e. the contribution of size-independent mortality sources to total mortality increases, given a constant life expectancy in habitat 2), the body size at habitat shift increases, so individuals have a delayed habitat shift with respect to the individuals exposed only to size-dependent mortality (fig. 2B; notice the same life expectancy at the beginning of the evolutionary trajectories in fig. 2D). By shifting habitat at a larger size, individuals increase their life expectancy in habitat 2 (fig. 2D).

In summary, the juveniles switch to the risky habitat 2 at a smaller body size when the risk is more size-dependent, despite that the risk would be avoidable by staying longer and growing in the nursery habitat to safety in size. This result is robust to a different size-dependent mortality function (Supp. Info. fig. S2, S3). In the following subsections, we explain the cause of this apparent evolutionary paradox.

### Increased size-selectivity in mortality in habitat 2 decreases growth potential in habitat 1

An analysis of the fitness components of the resident phenotype in the environment set by the resident population before and after the change in size-selectivity of mortality at evolutionary time 100 000 shows that an increase in size-selectivity in mortality in habitat 2 results in a slower growth rate and thus longer stay in habitat 1, higher survival and later maturation (black lines in fig. 3, left column). In contrast, this analysis reveals that a decrease in size-selectivity in mortality in habitat 2 results in a shorter stay in habitat 1 as a consequence of the increased growth rate in this habitat, lower survival and earlier maturation (black lines in fig. 3, right column).

Similarly, comparison of the resident and mutant phenotypes in the environment set by the resident after the change in size-selectivity of mortality at evolutionary time 100 000 shows that after an increase in size-selectivity in mortality, a smaller body size at habitat shift is selected for as it maximizes growth rate, leads to earlier maturation and increases fecundity at the expense of lower survival (dashed lines in fig. 3, left column). This analysis shows as well that after a decrease in size-selectivity in mortality a larger body size at the habitat shift is selected for as it maximizes survival at expense of slower growth and thus, later maturation (dashed lines in fig. 3, right column).

Fig. 3 shows that variation in size-selectivity in mortality in habitat 2 produces changes in the individual fitness components that are countered subsequently by selection. While we expected a direct effect of size-selectivity in mortality on survival, its effect on growth rate in the nursery habitat needs further explanation.

### By shaping population structure, size-selectivity in mortality influences growth potential in habitat 1

Size-selectivity in mortality in habitat 2 causes changes in the size-distribution of the population (the effects of size-dependent and size-independent mortality on habitat shift and population structure are further studied in Supp. Info. Fig. S4, S5). Because small individuals experience higher mortality rates than large individuals in habitat 2, adult density increases and juvenile density decreases in this habitat (fig. 4 left panel) when the size-selectivity in mortality is increased. This larger density of adults produces more offspring, which raises the density of juveniles in habitat 1. As a consequence of the increased density of juveniles in habitat 1, competition for food resources is stronger and thus growth rate is slower in this habitat. Given the adverse effects of density on growth potential, by advancing their shift to habitat 2 individuals can escape at an earlier age the reduced body growth they experience in habitat 1.

In contrast, a decreased size-selectivity in mortality in habitat 2 causes an increase in juvenile density and a decrease in adult density in this habitat (fig. 4 right panel). With a reduction in adult density the population birth rate decreases and thus the density of juveniles in habitat 1. Therefore, competition is relaxed and growth rate in this habitat increases. With a high growth potential in habitat 1, a later habitat shift enables individuals to increase their survival by postponing the shift to the riskier habitat 2.

In summary, we expected juveniles to switch to the risky habitat 2 at a larger body size when the risk is more size-dependent, because the risk would be avoidable by growing in the nursery habitat to safety in size. However, the opposite –a habitat shift at a smaller body size– was observed because the potential for growth in body size in the nursery habitat is low when the risk is more size-dependent in the risky habitat. Interestingly, this evolutionary response of reducing the body size at habitat shift when size-selectivity in the risky habitat increases is strong when total mortality in this habitat is low (or life expectancy is high) and becomes less strong as mortality increases (life expectancy decreases) (fig. S6). This is the consequence of stronger density dependence effects under low mortality than under high mortality conditions.

## Discussion

We have found an unexpected evolutionary response of the timing of a habitat to changes in size-selectivity in mortality. Our naïve expectation, based on an individual-level optimization, was that when size-selectivity in mortality in the rich feeding habitat increases, the body size at habitat shift would increase because delaying the habitat shift would cause individuals to benefit from increased survival in a larger part of their life cycle than when mortality is random across all size classes. In fact, this is the case when an analysis of fitness maximization of this life history trait is carried out for an isolated individual (fig. 5). In contrast, when accounting for density dependence in the ‘nursery’ habitat, the structured population model shows that the body size at habitat shift decreases with an increasing size-selectivity in mortality in the rich feeding habitat. This is the consequence of the effect that size-dependent and size-independent mortality in the risky habitat have on the population structure. Specifically, by changing the population structure, higher size-selectivity in mortality increases the density of juveniles in the ‘nursery’ habitat resulting in increased competition and, thus, triggering an earlier habitat shift.

**Figure 5.**
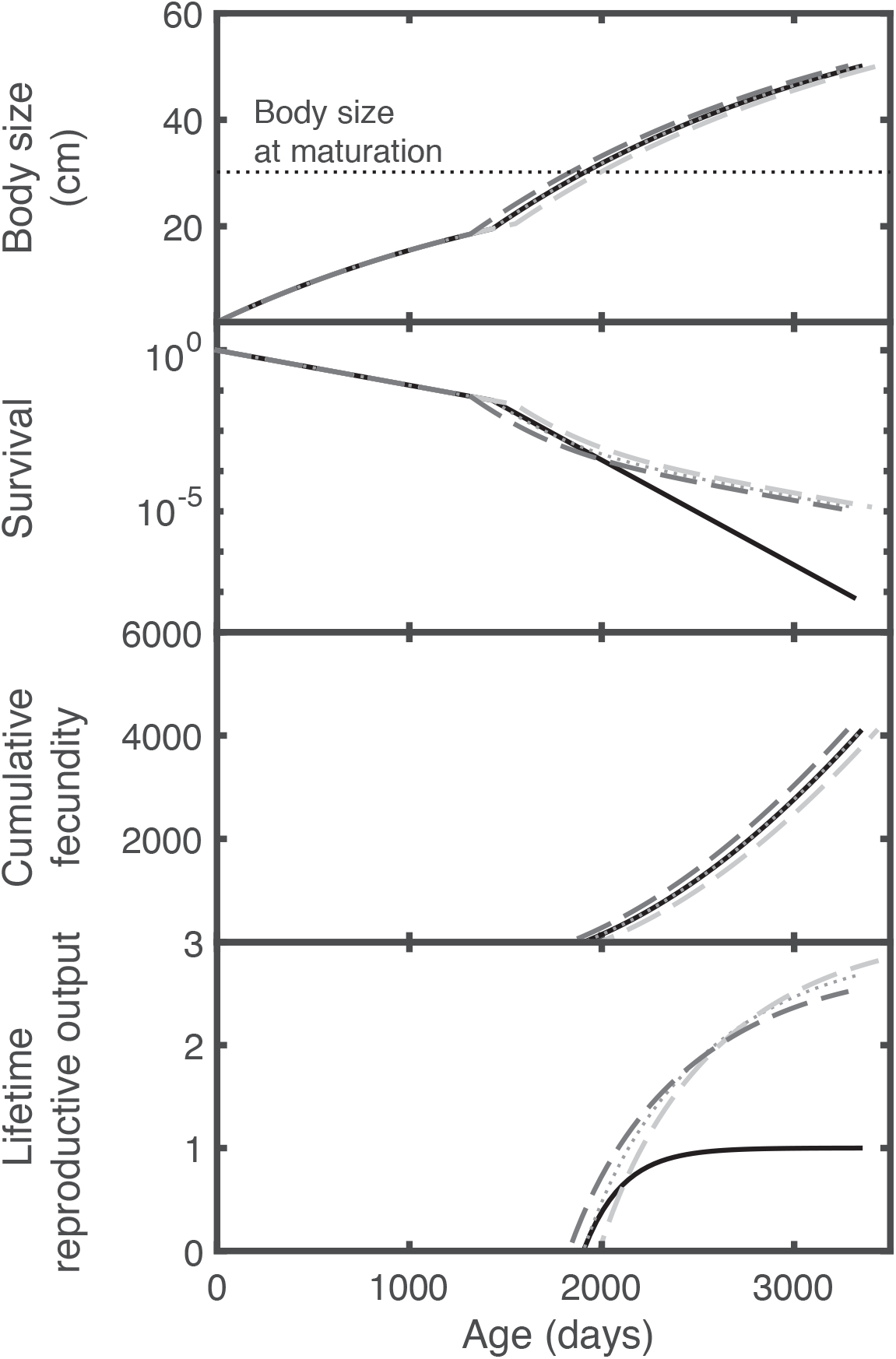
Individual growth, survival, cumulative fecundity and expected lifetime cumulative fecundity before (solid line) and after (dashed and dotted lines) size-selectivity in mortality increases in habitat 2 for an individual in isolation (no density-dependence in habitat 1). Initial phenotype (dotted line) shifts habitat at 19.5 cm, and novel phenotypes shift habitat at a body size 5% larger than the initial phenotype (light grey dashed lines) and at a body size 5% smaller than the initial phenotype (dark grey dashed lines). Size-dependent and size-independent mortality before (after) size-selectivity in mortality increases in the habitat 2 are as shown in table 1 for Pop. # 1 (Pop. # 3). A larger fitness is achieved by a novel phenotype with larger body size at habitat shift after the increase in size-selectivity in mortality if there is no density-dependence in the habitat 1 (in all cases individuals grow at the same rate: R_1_ is constant and equal to 0.457 g/L).

We have shown that mortality in the rich feeding habitat affects the optimal timing of the habitat shift not only because of its direct effect on survival but also through indirect effects on other fitness components such as growth. Werner and Gilliam (1984) have hypothesized before that the optimal timing of a habitat shift is determined by both the mortality and growth rate in the two habitats, and that the mortality rate is largely dependent on growth. Yet, the opposite effect that mortality influences growth by regulating the strength of density-dependence is a recent concept introduced by structured population theory: by relaxing competition, mortality affects and in particular promotes food-dependent processes such as growth and reproduction (de Roos et al. 2007). Furthermore, such effects have proven to shape population structure (i.e. size-distribution) and life history traits (Claessen et al. 2002; de Roos et al. 2007). In line with those findings for a population within a single habitat, we show that when there is a habitat shift, the nature of mortality in the rich feeding habitat has effects on growth, a food-dependent process, in the ‘nursery’ habitat.

Multiple studies have reported density, food availability and growth rate as determinants of the habitat shift in both experimentally manipulated as well as wild populations. For instance, experimental manipulations have shown that Brown trout is more likely to migrate (shift habitat) when growing slowly at high density but less likely to do so when density is low and growth rate is high (Olsson et al. 2006). This effect was proven to be mediated by food availability (Wysujack et al. 2009); and, like Brown trout, Arctic char (Nordeng 1983) and Atlantic salmon (Lans et al. 2011)are more likely to migrate at low food availability causing slow growth rate. Similarly, low food availability causing slow growth rate results in smaller size at metamorphosis than that of fast growers in amphibians (Alford and Harris 1988; Beachy, Surges, and Reyes 1999). In wild populations observations are similar: a long-term study of Atlantic salmon in the Simojoki river showed that the mean body size at smolting (habitat shift) was negatively correlated with density in the previous autumn (Jutila et al. 2006). Hence, high population densities depress food levels and thereby growth rates, which triggers an early habitat shift in different species with ontogenetic habitat shift. This effect that growth rate has on the optimal timing of the habitat shift is well known from an individual-level optimization perspective, however its connection with density-dependent processes resulting from feedbacks between the population and the individual life history are only recently starting to be explored. In this study, we showed the intimate linkage between population structure and the optimal timing of the habitat shift as they have reciprocal effects on each other. Given the multiple and dramatic consequences that population structure and habitat shifts have independently on communities and ecosystems (de Roos and Persson, 2002; Schreiber and Rudolf, 2008), the implications of interactions between them need to be studied in future research.

In this study we have focused on size-dependent predation mortality in the rich feeding habitat whose size-selectivity implies a negative relation between mortality risk and body size. However, a positive relation between mortality risk and body size is also common. For instance, in general, fishing mortality imposed on exploited fish populations is higher for larger individuals. Fish stocks, therefore, may experience strong positive size-selective mortalities. Since adult biomass increases when the proportion of negative size-selective predation mortality increases as mortality cause, size-dependent fishing mortality targeting mainly large individuals would reduce the adult biomass; hence, it cancels out the effect on population structure that size-dependent predation mortality has. As a consequence, in a population with size-dependent predation mortality as main source of natural mortality, size-dependent fishing mortality would reduce density and relax competition in the ‘nursery’ habitat. Consequently, the combination of negative size-selective predation mortality and positive size-selective fishing mortality would promote a habitat shift at larger body size. Indeed, Atlantic salmon in the Baltic sea has experienced a drop in the fishing effort in the last decades with concurrent higher density of individuals in the ‘nursery’ Simojoki river resulting in smaller mean sizes at the habitat shift (Jutila et al. 2006). Survival in early stages after the habitat shift are correlated with body size at the shift; therefore, Simojoki river Atlantic salmon is experiencing lower survival after the habitat shift. This suggests that some size-dependent fishing mortality may actually increase survival after habitat shift and perhaps enhance the fishing yield. Further research is necessary to determine the effects of fishing mortality on the optimal timing of the habitat shift.

For simplicity, in this study we have examined the evolution of the timing of a habitat shift while other life history traits are kept fixed. However, other life history traits are likely to evolve as well in response to changing fitness components like survival and growth. In particular, individuals may mature at smaller or larger size depending on growth opportunities and mortality risk. Fish often attain sexual maturation when growth rate reduces (near the asymptotic body size) (Jonsson and Jonsson 1993). Since the habitat shift enables individuals to access a rich feeding habitat and thus rapid growth, the timing of the habitat shift may influence the optimal timing of maturation. For instance, Brown trout moving from ‘nursery’ streams to lakes for feeding delays maturity in comparison with stream residents for one or more years (Jonsson 1989). In future research, it would be therefore interesting to study jointly the evolution of the timing of the habitat shift and the timing of sexual maturation. Given that both traits influence population structure, such work will contribute to a deeper understanding of the linkage between life history trait evolution and density-dependent effects.

We consider the evolution of the body size at habitat shift in a monomorphic population in our model. However, in the wild, organisms with habitat shift, like salmonids and coral reef fish, often show variation in body size at habitat shift within a population. An extension of this model may include plasticity in this life history trait. For instance, adopting a reaction norm approach in which the probability to shift habitat is a function of the body size. Such an approach has been previously used to asses the evolutionary outcome of different mortality sources in the absence of density-dependent effects (Thériault et al. 2008). An extension of this approach to account for density-dependence would be possible using the Physiologically Structured Population Model framework used in this study (de Roos 1997).

Survival and growth rate have long been recognized as the traits to optimize when shifting habitats. In addition, a growing body of theoretical work and experimental evidence shows that survival and growth rate are interdependent and interact through feedbacks between the population and the individual life history. Despite this, the analysis of the optimal timing of a habitat shift, as well as other life history traits, has been traditionally carried out in a context of individual optimization in isolation. Our results demonstrate the strong mutual influence that population structure and optimal timing of a habitat shift have on each other. This highlights the need for integrating ecological interactions in the study of optimal life history traits. If we are to understand the evolution of life histories the integration of ecological interactions in the evolutionary analysis is certainly required.

