## Supplementary material for "Density-dependent effects of mortality on the optimal body size of a habitat shift: Why smaller is better despite increased mortality risk?"

This supplementary note presents the following sections:

1. Size-structured population model
2. The evolutionary model and canonical equation
3. Model parameters
4. Robustness of results
5. Effects of size-dependent and size-independent mortality on habitat shift and population structure

### 1. Size-structured population model

The population is structured by size and follows continuous-time dynamics. Derivation of the structured population follows from the individual dynamics description (de Roos et al. 1990; de Roos 1997).

The dynamics of the size-structured population in the two habitats are described by the following partial differential equations:

$$\frac{\partial c_1(t,l)}{\partial t} + \frac{\partial(\gamma(f_1,l)\, c_1(t,l))}{\partial l} = -\, \mu_1 c_1(t,l) \tag{S1.1}$$

$$\frac{\partial c_2(t,l)}{\partial t} + \frac{\partial(\gamma(f_2,l)\, c_2(t,l))}{\partial l} = -\, (\mu_{2\,b} + \mu_{2\,p}(l))c_2(t,l) \tag{S1.2}$$

$$\gamma(f_2,l_s)c_2(t,l_s) = \gamma(f_1,l_s)c_1(t,l_s) \tag{S1.3}$$

$$\gamma(f_1,l_0)c_1(t,l_0) = \begin{cases} \int_{l_m}^{l_s} \beta(f_1,l)\, c_1(t,l)\, dl + \int_{l_s}^{\infty} \beta(f_2,l)\, c_2(t,l)\, dl & \text{if } l_m < l_s \\ \int_{l_m}^{\infty} \beta(f_2,l)\, c_2(t,l)\, dl & \text{otherwise} \end{cases} \tag{S1.4}$$

where, $c_1(t,l)$ and $c_2(t,l)$ are the body size-distributions of individuals in habitat 1 and habitat 2, respectively.

Equations (3) and (4) provide the boundary condition at the switching size $l_s$ from habitat 1 to habitat 2 and at the birth size $l_0$, respectively. The right-hand side of the latter boundary condition corresponds to the total population birth rate.

The food resource density in habitat 1 increases by renewal as explained in eq. 1 and decreases by consumption following

$$\frac{dR_1}{dt} = \rho(R_{1\,\max} - R_1) - \frac{R_1}{K + R_1}\frac{j'_a}{\zeta_a}\int_{l_0}^{l_s} l^2 c_1(t,l)dl \tag{S1.5}$$

in which, $\zeta_a$ is the efficiency with which ingested food is assimilated.

The total biomass of juveniles and adults in both habitats together is calculated with the integral expressions

$$\begin{cases} \displaystyle\int_{l_0}^{l_m} v\,(\delta\, l)^3\, c_1(t,l)\, dl & \text{if } l_m < l_s \\ \displaystyle\int_{l_0}^{l_s} v\,(\delta\, l)^3\, c_1(t,l)\, dl + \int_{l_s}^{l_m} v\,(\delta\, l)^3\, c_2(t,l)\, dl & \text{otherwise} \end{cases} \tag{S1.6}$$

and

$$\begin{cases} \displaystyle\int_{l_m}^{l_s} v\,(\delta\, l)^3\, c_1(t,l)\, dl + \int_{l_s}^{\infty} v\,(\delta\, l)^3\, c_2(t,l)\, dl & \text{if } l_m < l_s \\ \displaystyle\int_{l_m}^{\infty} v\,(\delta\, l)^3\, c_2(t,l)\, dl & \text{otherwise} \end{cases}, \tag{S1.7}$$

respectively.

### 2. Evolutionary model and canonical equation

We used the canonical equation of adaptive dynamics to describe the evolutionary rate of change in the body size at habitat shift $l_s$ (Dieckmann and Law 1996; Durinx et al. 2008). The evolutionary rate of change of the trait is proportional to the selection gradient and depends on the production rate of mutants, and their establishment chance,

$$\frac{dl_s}{d\tau} = \frac{T_f}{T_s} \frac{\hat{n}\,\vartheta}{\sigma^2} \frac{\partial s(l'_s|l_s)}{\partial l'_s}\bigg| l'_s = l_s \qquad \text{(S2.1)}$$

where $\tau$ spans the evolutionary timescale. In this equation $T_f$ is the average age at giving birth, $T_s$ is the expected lifespan of an individual, $\hat{n}$ is the size of the population in equilibrium, $\vartheta$ is the mutation probability per birth event, $\sigma^2$ is the variance of the offspring trait distribution and $\frac{\partial s(l'_s|l_s)}{\partial l'_s}\Big| l'_s = l_s$ is the selection gradient. Furthermore, the ratio $\hat{n}/T_s$ equals the total birth rate $\hat{b}$ of the population in equilibrium. According to Durinx et al. (2008) the relation between $R_0$ and the invasion fitness $s(l'_s|l_s)$ is given by

$$s(l'_s|l_s) = \frac{log\big(R_0(l'_s|l_s)\big)}{T_f} \qquad \text{(S2.2)}$$

Such that the canonical equation for the body size at habitat shift $l_s$ can be written as:

$$\frac{dl_s}{d\tau} = C\,\hat{b}\,\frac{\partial R_0(l'_s|l_s)}{\partial l'_s}\bigg| l'_s = l_s \qquad \text{(S2.3)}$$

given that $R_0(l_s'|l_s') = 1$. In the above equation the constant $C = \vartheta/\sigma^2$ scales the evolutionary rates of the evolutionary trajectories. We chose the value of $C$ equal to 1 because the evolutionary time units are considered arbitrary. The PSPManalysis (de Roos 2018) package implements the form (S2.3) of the canonical equation with $C = 1$ to study evolutionary dynamics by numerically calculating the values of $R_0$, $\hat{b}$ and the selection gradient $\partial R_0(l_s'|l_s)/\partial l_s'$ from the life history functions for growth, mortality and reproduction.

### 3. Model parameters

Table S1. Parameter values

| Description | Symbol | Value | Unit | References |
|---|---|---|---|---|
| Resource in the 'nursery' habitat | | | | |
| Resource growth rate | $\rho$ | 0.01 | day$^{-1}$ | |
| Maximum resource density | $R_{1\,max}$ | 4 | g m$^{-3}$ | |
| Population with habitat shift | | | | |
| Half saturation resource density | $K$ | 1 | g m$^{-3}$ | |
| Feeding level in the habitat 2 | $f_2$ | 0.6 | - | |
| Fraction of assimilation flux to structural mass and maintenance | $\kappa$ | 0.8 | | (Jager et al. 2013) |
| Maximum area-specific assimilation rate | $j_a$ | 0.0572* | g g$^{-2/3}$ day$^{-1}$ | Calculated with method of (Jager et al. 2013) from regressions of (Koskela et al. 1997) |
| Mass-specific maintenance cost | $j_m$ | 0.0019* | g g$^{-1}$ day$^{-1}$ | Calculated with method of (Jager et al. 2013) from regressions of (Koskela et al. 1997) |
| Yield of structural mass on assimilates | $\zeta_a$ | 1 | g g$^{-1}$ | (Jager et al. 2013)(Jager et al. 2013)(Jager et al. 2013)(Jager et al. 2013)(Jager et al. 2013) |
| Yield of structural mass on assimilates | $\zeta_W$ | 0.8 | g g$^{-1}$ | (Jager et al. 2013) |
| Yield of egg buffer on assimilates | $\zeta_e$ | 0.8 | g g$^{-1}$ | (Jager et al. 2013) |
| Shape coefficient factor | $\delta$ | 0.21 | - | (Pecquerie et al. 2011) |
| Density of structural mass | $v$ | 1 | g cm$^{-3}$ | (Jager et al. 2013) |
| Body size of a newborn | $l_0$ | 2 | cm | (Gilbey et al. 2009) |
| Body size at the habitat shift | $l_s$ | varied (evolving) | cm | |
| Body size at maturation | $l_m$ | 30 | cm | |
| Maximum vulnerable-to-predation body size** | $l_v$ | 50 | cm | |
| Maximum size-dependent mortality** | $\varepsilon$ | varied | day$^{-1}$ | |
| Scaling coefficient of size-dependent mortality*** | $c$ | varied | day$^{-1}$ | |
| Exponent of size-dependent mortality*** | $b$ | 0.75 | - | (Jørgensen and Holt 2013) |
| Egg survival | $\sigma$ | 0.5 | - | (Bley & Moring, 1988) |
| Mortality rate in the habitat 1 | $\mu_1$ | 0.002 | day$^{-1}$ | (Bley & Moring, 1988) |
| Size-independent mortality rate in the habitat 2 | $\mu_{2\,b}$ | varied | day$^{-1}$ | |

*The rate constant ($j_a$, $j_m$) values include a temperature correction following the Arrhenius relationship for a temperature of 10 °C.

**Parameters used only when size-dependent mortality is a sigmoid function of the body size (equation 11).

***Parameters used only when size-dependent mortality is an exponential function of the body size (equation 12).

### 4. Robustness of results

In this section, we study the robustness of our results by assuming a different function for size-dependent mortality in habitat 2. Specifically, we assume the size-dependent mortality to be an exponential function of body size (equation 12 in Methods section, fig S1).

To unravel the effect of the size-selectivity of mortality, we follow a similar procedure as described in the methods section for the sigmoid function of size-dependent mortality. To ensure that overall mortality in habitat 2 is kept constant when increasing size-selectivity, the contribution from size-dependent mortality sources is increased and simultaneously size-independent mortality is decreased. In contrast to the procedure in the methods section of the main text we used $c$, instead of $\varepsilon$, as the parameter to define the magnitude of the size-dependent mortality. More specifically, we find the ESS value for the body size at habitat shift when there is only size-independent mortality $\mu_{2\,b}$ equal to 0.006 day$^{-1}$ in habitat 2 and adopt this as our starting, reference population (body size at habitat shift = 19.5 cm). A size-independent mortality $\mu_{2\,b}$ equal to 0.006 day$^{-1}$ implies that an individual has an expected lifetime of 167 days in habitat 2 from the moment it shifts habitats. Adopting this ESS body size for the resident phenotype in case it only experiences the size-independent mortality $\mu_{2\,b}$ of 0.006 day$^{-1}$, we infer two combinations of size-independent mortality $\mu_{2\,b}$ and the scaling coefficient of size-dependent mortality $c$ that also result in a life expectancy of 167 days in habitat 2 (Pop. # 2 and 3 in table S2). We subsequently start with a resident population that is characterized by the ESS value for body size to shift habitat while experiencing a size-independent mortality equal to 0.006 day$^{-1}$ in habitat 2 and assess its evolutionary response when the mortality changes to one of the inferred combinations of size-independent mortality

$\mu_{2\,b}$ and the scaling coefficient of size-dependent mortality $c$ that also lead to an average life expectancy of 167 days in habitat 2 (fig. S2A). We perform a similar analysis while decreasing the size-selectivity in mortality in habitat 2: We adopt as our starting, reference population, one characterized by the ESS value for the body size to shift habitat when size-dependent mortality is the main source of mortality with a scaling coefficient $c$ of 0.02 day$^{-1}$ in habitat 2 (body size to shift habitat = 19.5 cm; a very low size-independent mortality equal to 0.0001 day$^{-1}$ in habitat 2 is introduced to avoid that individuals with a large body size become immortal). We identify again two other combinations of the scaling coefficient of size-dependent mortality $c$ and the size-independent mortality $\mu_{2\,b}$ that result in the same life expectancy in the habitat 2 (Pop. # 5 and 6 in table S2) as the size-dependent mortality with scaling coefficient $c$ of 0.02 day$^{-1}$ for individuals shifting habitat at the ESS body size and study the evolutionary response of the starting resident population to a change to the two inferred mortality schedules (fig. S2B).

For the four combinations of size-dependent and size-independent mortalities mentioned above we use the PSPManalysis software package to compute the evolutionary trajectories of the body size at the habitat shift (evolutionary time 100 000 to 400 000 in fig. 2) as predicted by the canonical equation of adaptive dynamics. In addition, we assess the effect of the variation in size-selectivity in mortality on the size distribution of the population (fig. S3).

Table S2. Mortality parameters used in Fig. S2 and S3.

| **Pop. #** | **Body size at habitat shift[a] (cm)** | **Life expectancy in habitat 2 (days)** | **Scaling coeff. size-dependent mortality $c$** | **Contribution of size-dependent mortality [b]** | **Size-independent mortality $\mu_{2\,b}$** | **Contribution of size-independent mortality [b]** |
|---|---|---|---|---|---|---|
| | Increasing size-selectivity in mortality in habitat 2 | | | | | |
| 1[d] | 19.5 | 167 | 0 | 0% | 0.006 | 100% |
| 2 | 19.5 | 167 | 0.0065 | 13.1% | 0.004 | 86.9% |

| | | | | | | |
|---|---|---|---|---|---|---|
| 3 | 19.5 | 167 | 0.0131 | 36.5% | 0.002 | 63.5% |
| | Decreasing size-selectivity in mortality in habitat 2 | | | | | |
| 4[d] | 19.5 | 161 | 0.02 | 94.1% | 0.0001 | 5.9% |
| 5 | 19.5 | 161 | 0.015 | 44.6% | 0.0016 | 55.4% |
| 6 | 19.5 | 161 | 0.01 | 22.5% | 0.0031 | 77.5% |

[a] Body size at habitat shift is ESS before change in size-selectivity in mortality.

[b] Contribution of each source of mortality to total mortality in habitat 2.

[d] Starting, reference population.

Figure S2 shows that when size-selectivity in mortality increases in habitat 2, the body size at habitat shift decreases (panel A), whereas it increases when size-selectivity in mortality decreases (panel B). This is the same qualitative result when size-dependent mortality is a sigmoid function of body size (fig. 2). The evolutionary response towards reduced body size at habitat shift when size-selectivity increases is related to an increase in density of juveniles in habitat 1 (fig. S3 left panel). The size-distribution of the population shows the same qualitative pattern when size-selectivity increases or decreases regardless of the function that defines size-dependent mortality. Qualitatively, the main result in our paper is hence robust despite that the different shapes of the size-dependent mortality functions cause the size-distributions of the population to be quantitatively different (fig. 4 vs. fig. S3).

### 5. Effects of size-dependent and size-independent mortality on habitat shift and population structure

In this section we studied the effect of varying size-dependent and size-independent mortality on the body size at the habitat shift and its effects on population structure. To do so, we examine the effect of increasing size-dependent mortality only when there is very low size-independent mortality in the habitat 2 ($\mu_{2\,b} = 0.00001$) on the body size at habitat shift (fig. S4 left panel). Conversely, we study the effect of increasing size-independent mortality only when there is no size-dependent mortality in the habitat 2 on the body size at habitat shift (fig. S4 right panel). Subsequently, we evaluate the population structure (i.e. biomass density of juveniles in habitat 1 and in habitat 2, and of adults) occurring if individuals shift habitat at the ESSs when size-dependent and size-independent mortality vary as indicated above.

Figure S4 shows that with increasing mortality in habitat 2, regardless of its nature (size-dependent or size-independent), body size at the habitat shift increases. However, as body size at the habitat shift increases its effects on the population structure vary depending on the nature of the mortality in habitat 2 (fig. S5). When mortality in habitat 2 is caused by size-dependent mortality, small individuals experience higher mortality rates than large individuals in this habitat. In contrast, when mortality in habitat 2 is caused by size-independent mortality, all individuals in this habitat experience similar mortality rates regardless of their body size. As a consequence, when size-dependent mortality is the only source of mortality in habitat 2, juveniles in this habitat are less and adults are more abundant compared to a population experiencing only size-independent mortality in the same habitat and shifting at the same optimal body size (fig. S5). A larger adult population produces more offspring and thus results in a larger juvenile biomass density in habitat 1. Because a large biomass of juveniles in habitat 1 consumes and depletes the food resource in this habitat to lower density (fig. S5, bottom plot), individuals in this habitat have low feeding rate and thus experience slower growth rates.

de Roos, A. M. 2018. PSPManalysis: A package for numerical analysis of physiologically structured population models.

de Roos, A. M., J. A. J. Metz, E. Evers, and A. Leipoldt. 1990. A size dependent predator-prey interaction: who pursues whom? Journal of Mathematical Biology 28:609–643.

Dieckmann, U., and R. Law. 1996. The dynamical theory of coevolution: a derivation from stochastic ecological processes. Journal of Mathematical Biology 34:579–612.

Durinx, M., J. A. J. (Hans) Metz, and G. Meszéna. 2008. Adaptive dynamics for physiologically structured population models. Journal of Mathematical Biology 56:673–742.

Gilbey, J., E. Cauwelier, C. S. Jones,  a. McLay, L. R. Noble, and E. Verspoor. 2009. Size-dependent growth of individual Atlantic salmon Salmo salar alevins from hatch to first feeding. Journal of Fish Biology 75:2820–2831.

Jager, T., B. T. Martin, and E. I. Zimmer. 2013. DEBkiss or the quest for the simplest generic model of animal life history. Journal of Theoretical Biology 328:9–18. Elsevier.

Jørgensen, C., and R. E. Holt. 2013. Natural mortality: Its ecology, how it shapes fish life histories, and why it may be increased by fishing. Journal of Sea Research 75:8–18. Elsevier B.V.

Koskela, J., J. Pirhonen, and M. Jobling. 1997. Feed intake, growth rate and body composition of juvenile Baltic salmon exposed to different constant temperatures. Aquaculture international 5:351–360.

Pecquerie, L., L. R. Johnson, S. A. L. M. Kooijman, and R. M. Nisbet. 2011. Analyzing variations in life-history traits of Pacific salmon in the context of Dynamic Energy Budget (DEB) theory. Journal of Sea Research 66:424–433.

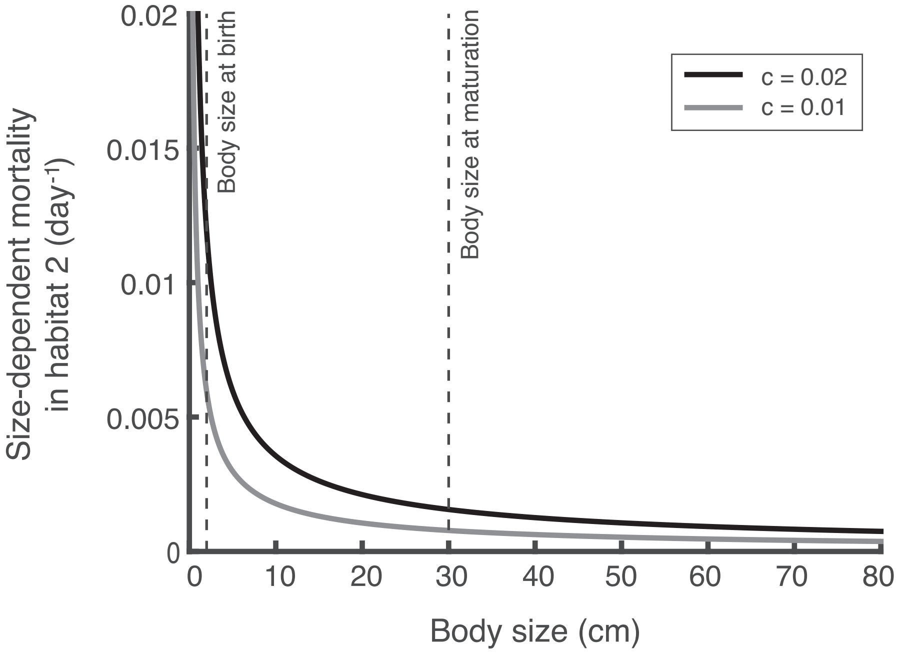


Figure S1. Size-dependent mortality in habitat 2 following an exponential function (Size-dependent mortality = c * (Body size)$^{-b}$ ; b = 0.75).

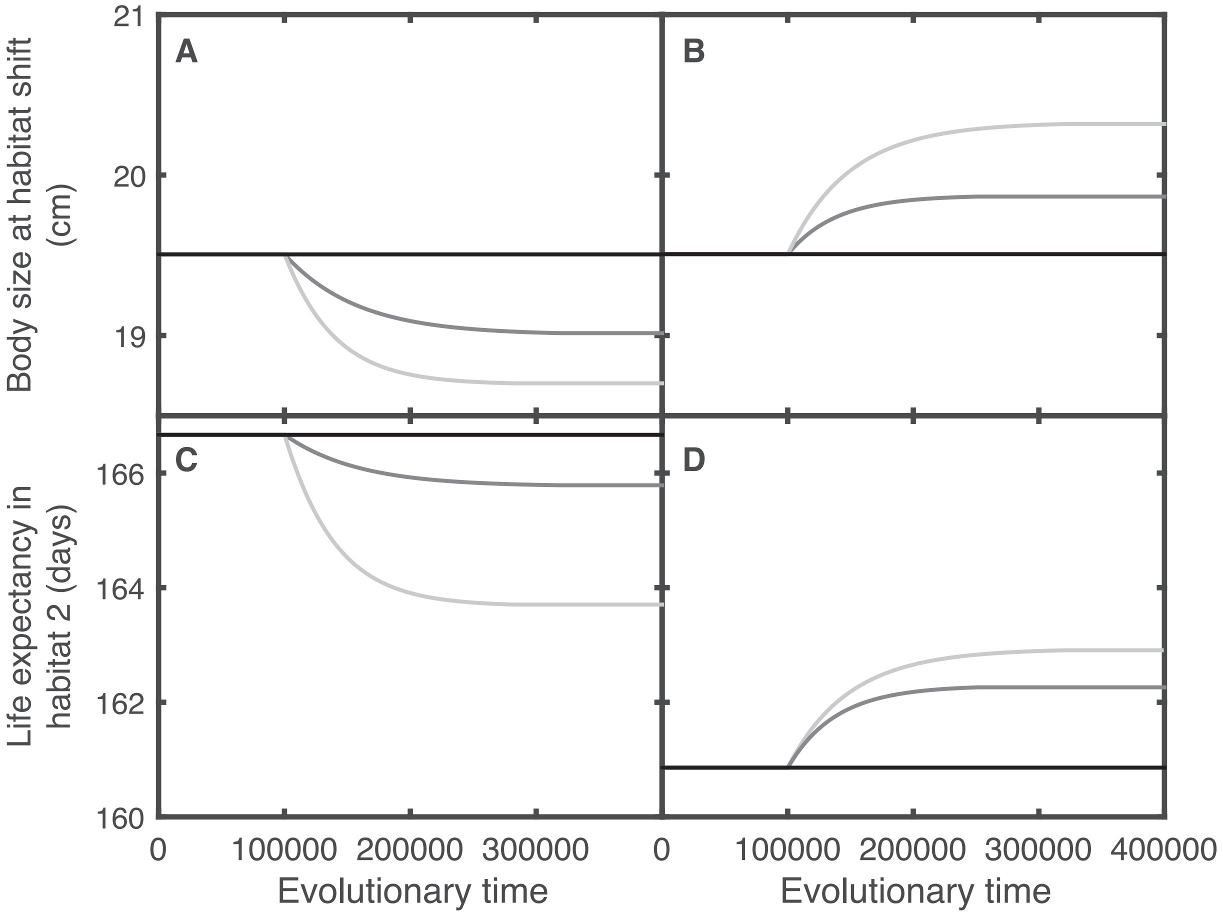


Figure S2. Effects of increased (A, C) and decreased (B, D) size-selectivity in mortality in habitat 2 on the optimal timing of the habitat shift (A, B) and life expectancy in habitat 2 (C, D) when size-dependent mortality in habitat 2 is an exponential function of the body size (fig. S1). When size-selectivity increases (A, C), evolutionary dynamics starts from the ESS value for the body size at habitat shift (19.5 cm) when individuals experience only size-independent mortality 0.006 $day^{-1}$ until time 100 000. At this time, the change in size-selectivity in mortality in habitat 2 occurs (changes in size-dependent and size-independent mortality according to table S2: Pop. # 1 corresponds to the black line, 2 to the dark grey line and 3 to the light grey line). When size-selectivity decreases (B, D), evolutionary dynamics starts from the ESS value for the body size at habitat shift (19.5 cm) when individuals experience mainly size-dependent mortality with a scaling coefficient c = 0.02 $day^{-1}$ until time 100 000. At this time, the change in size-selectivity in mortality in habitat 2 occurs (changes in size-dependent and size-independent mortality according to table S2: Pop. # 4 corresponds to the black line, 5 to the dark grey line and 6 to the light grey line). Other parameter values as in table S1.

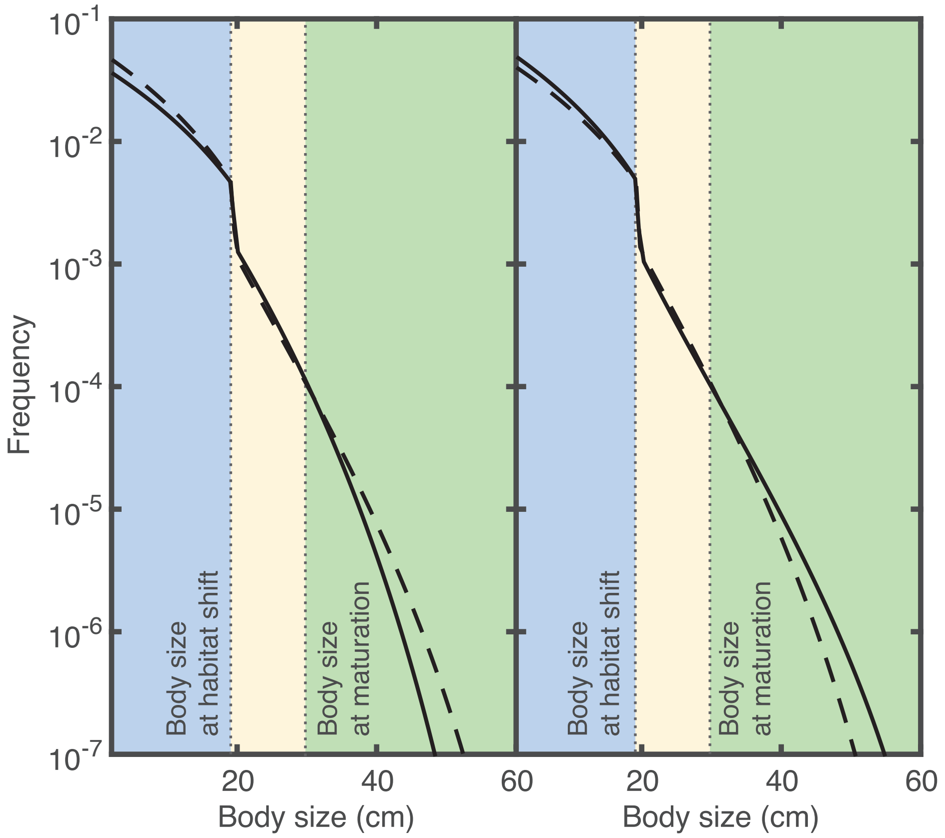


Figure S3. Size-distribution of the resident population just before (solid lines) and immediately after (dashed lines) size-selectivity in mortality increases (left column) and decreases (right column) in habitat 2. Individuals shift habitat at a body size of 19.5 cm. Juveniles in habitat 1 (blue region) and in habitat 2 (yellow region), and adults (green region) are shown. Size-dependent and size-independent mortality before size-selectivity in mortality increases and decreases are as shown in table S2 for Pop. # 1 and 4, respectively; whereas after the increase and decrease they are as shown in table S2 for Pop. # 3 and 6, respectively. Other parameter values as in table S1.

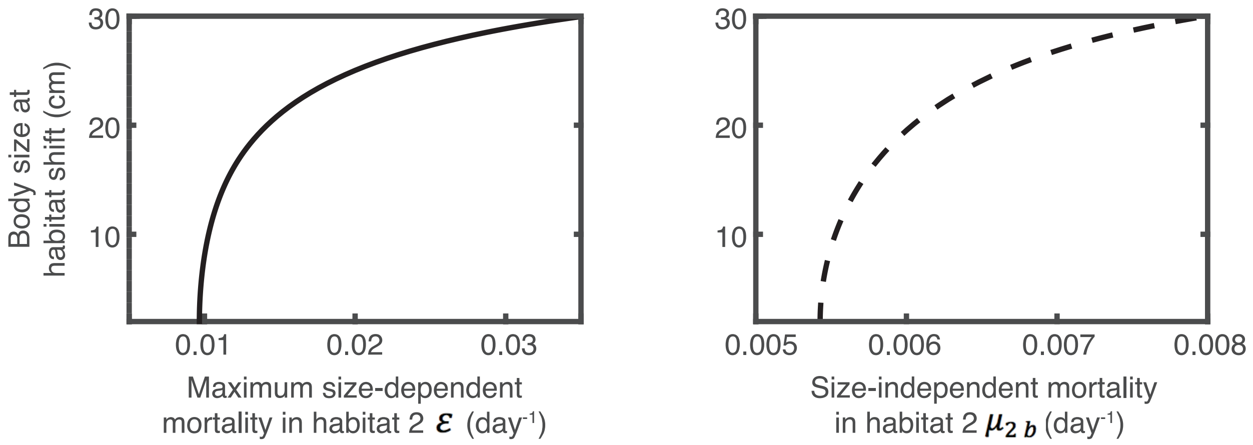


Figure S4. Body size at which individuals shift habitats at the ESS as a function of the maximum size-dependent mortality (left) and size-independent mortality in habitat 2 (right). Size-independent mortality is 0.00001 day$^{-1}$ when maximum size-dependent mortality varies and maximum size-dependent mortality is 0 days$^{-1}$ when size-independent mortality varies. Other parameter values are as shown in table S1.

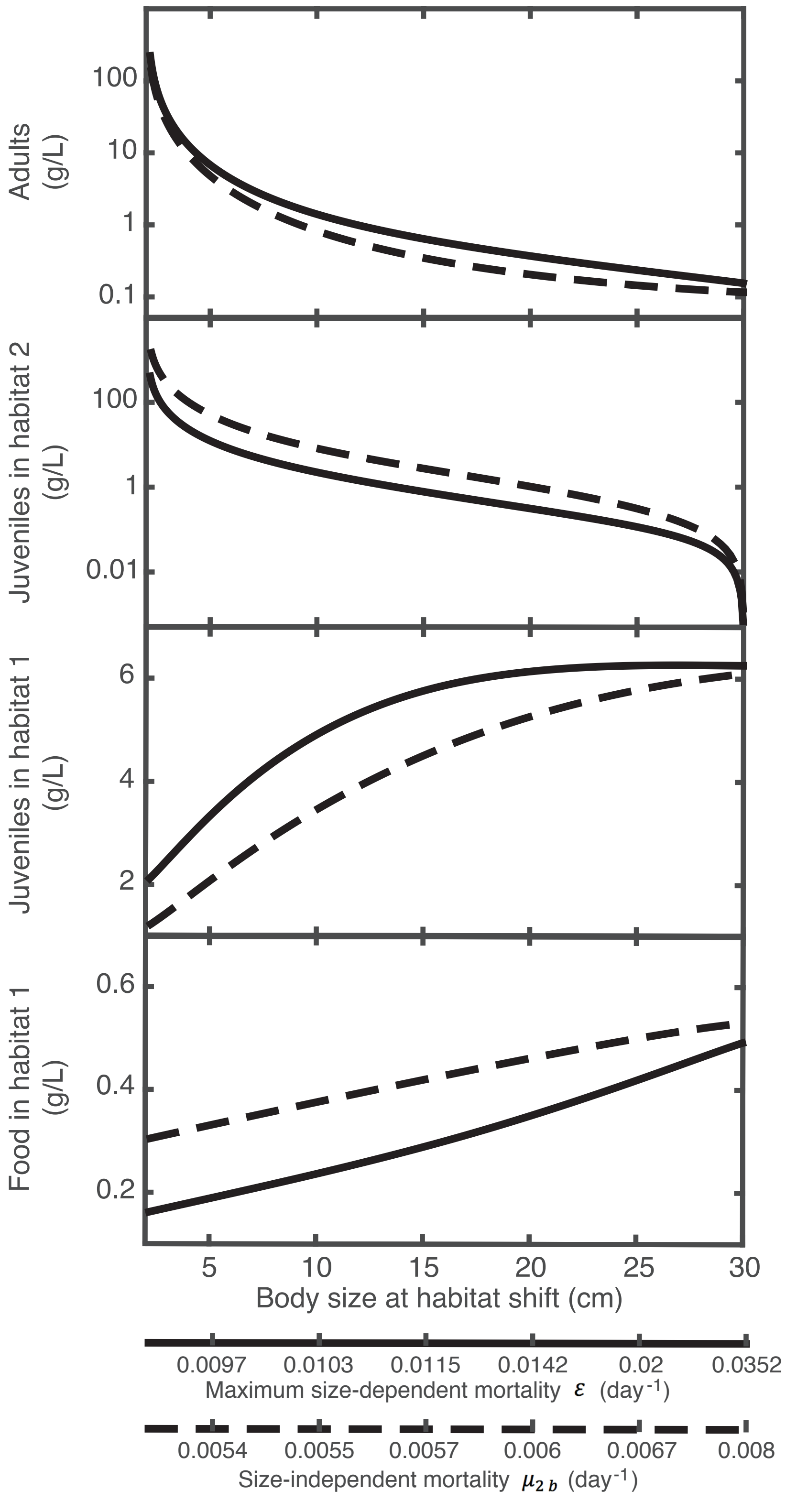


Figure S5. Adult biomass density, juvenile biomass density in habitat 1 and habitat 2 and food biomass density in habitat 1 as a function of the ESS value of the body size at the habitat shift when these changes in the ESS value of body size are due to variation in maximum size-dependent mortality in habitat 2 (solid line and x-axis; size-independent mortality is 1E-5) or to variation in size-independent mortality in the same habitat (dashed line and x-axis; no size-dependent mortality) as shown in figure S4. Other parameter values as in table S1.

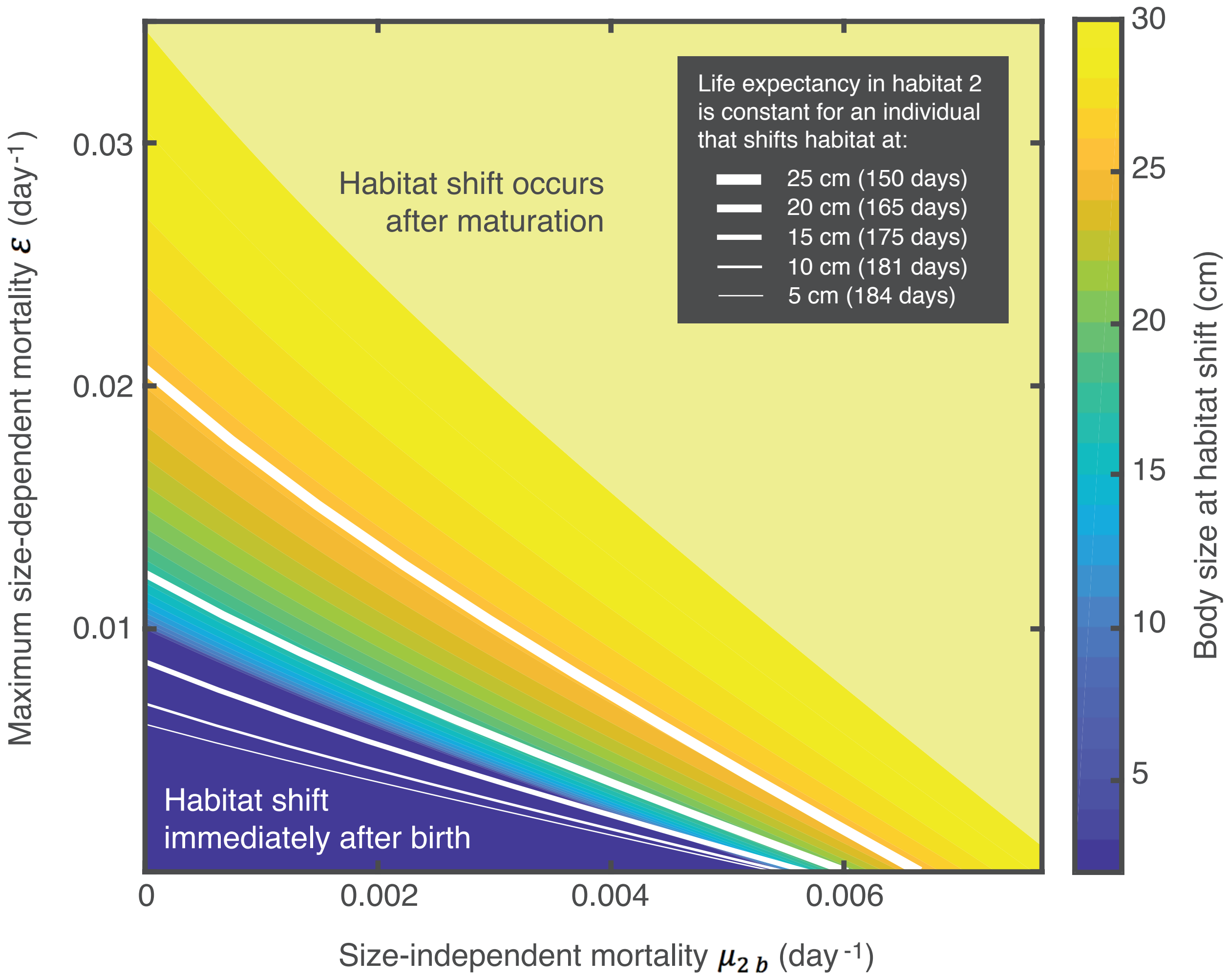


Figure S6. Body size at which individuals shift habitat (colorbar) at the ESSs as a function of the maximum size-dependent mortality (vertical axis) and the size-independent mortality (horizontal axis) in habitat 2. Isoclines (white lines) show the combinations of size-dependent and size-independent mortality at which the life expectancy in habitat 2 (in parentheses) is constant for an individual shifting habitat at a body size of 5, 10, 15, 20 and 25 cm. These body size values correspond to the ESSs when individuals only experience size-independent mortality and this mortality is equal to the value at the intersection of the isoclines with the horizontal axes.
